# The transcription factor IscR promotes *Yersinia* type III secretion system activity by antagonizing the repressive H-NS-YmoA histone-like protein complex

**DOI:** 10.1101/2021.10.26.466021

**Authors:** David Balderas, Pablo Alvarez, Mané Ohanyan, Erin Mettert, Natasha Tanner, Patricia J. Kiley, Victoria Auerbuch

## Abstract

The type III secretion system (T3SS) is a appendage used by many bacterial pathogens, such as pathogenic *Yersinia*, to subvert host defenses. However, because the T3SS is energetically costly and immunogenic, it must be tightly regulated in response to environmental cues to enable survival in the host. Here we show that expression of the *Yersinia* Ysc T3SS master regulator, LcrF, is orchestrated by the opposing activities of the repressive YmoA/H-NS histone-like protein complex and induction by the iron and oxygen-regulated IscR transcription factor. Although IscR has been shown to bind the *lcrF* promoter and is required for *in vivo* expression of *lcrF*, in this study we show IscR alone fails to enhance *lcrF* transcription *in vitro*. Rather, we find that in a *ymoA* mutant, IscR is no longer required for LcrF expression or T3SS activity. Additionally, a mutation in YmoA that prevents H-NS binding (*ymoA*^D43N^) rescues the T3SS defect of a Δ*iscR* mutant, suggesting that a YmoA/H-NS complex is needed for this repressive activity. Furthermore, chromatin immunoprecipitation analysis revealed that H-NS is enriched at the *lcrF* promoter at environmental temperatures, while IscR is enriched at this promoter at mammalian body temperature under aerobic conditions. Importantly, CRISPRi knockdown of H-NS leads to increased *lcrF* transcription. Collectively, our data suggest that as IscR levels rise with iron limitation and oxidative stress, conditions *Yersinia* experiences during extraintestinal infection, IscR antagonizes YmoA/H-NS-mediated repression of *lcrF* transcription to drive T3SS activity and manipulate host defense mechanisms.

**Author Summary:** Facultative pathogens must silence virulence gene expression during growth in the environment, while retaining the ability to upregulate these genes upon infection of a host. H-NS is an architectural DNA binding protein proposed to silence horizontally acquired genes, regulating virulence genes in a number of pathogens. Indeed, H-NS was predicted to regulate plasmid-encoded virulence genes in pathogenic *Yersinia*. However, *Yersinia* H-NS is reported to be essential, complicating testing of this model. We used chromatin immunoprecipitation and inducible CRISPRi knockdown to show that H-NS binds to the promoter of a critical plasmid-encoded virulence gene, silencing its expression. Importantly, under conditions that mimic *Yersinia* infection of a mammalian host, the transcriptional regulator IscR displaces H-NS to drive virulence factor expression.

## Introduction

Virulence factors are critical components that allow pathogens to establish or sustain infections within a given host. One common bacterial virulence factor is a needle-like apparatus, known as the type III secretion system (T3SS) (1,2). Enteropathogenic *Yersinia pseudotuberculosis* is one of three human pathogenic *Yersinia* spp. that use the T3SS to inject effector proteins into host cells that dampen host immune responses, facilitating extracellular growth (3–6). Members of human pathogenic *Yersinia* spp. include *Yersinia pestis*, the causative agent of plague, and the enteropathogens *Yersinia enterocolitica* and *Yersinia pseudotuberculosis*. While the T3SS is critical for infection, this apparatus appears to be metabolically burdensome since constitutive expression of the T3SS leads to growth arrest (7,8). In addition, the Ysc T3SS is associated with pathogen-associated molecular patterns (PAMPs) recognized by several innate immune receptors, and some of these T3SS-associated PAMPS have evolved under selective evolutionary pressure by the ensuing immune response (5,9). Without tight regulation of T3SS expression and deployment, these metabolic and immunological burdens would decrease the chance of *Yersinia* survival in the host.

The Ysc T3SS is encoded on a 70 kb <u>p</u>lasmid for *<u>Y</u>ersinia* <u>V</u>irulence, known as pYV or pCD1 (10). Transcriptional regulation of T3SS genes is maintained by a master regulator called LcrF/VirF (11–14). LcrF itself is also encoded on pYV, within the *yscW-lcrF* operon, and is highly conserved among all three human pathogenic *Yersinia* spp. LcrF is part of a larger family of AraC-like transcriptional regulators, and orthologs exist in other T3SS-encoding pathogens, such as ExsA in the nosocomial pathogen *Pseudomonas aeruginosa* (15). The *yscW-lcrF* operon is regulated at various stages in response to different environmental stimuli, including temperature, oxygen, and iron availability (16,17). For example, an RNA thermometer blocks the ribosome binding site of *lcrF* at room temperature, but melts at mammalian body temperature, allowing *lcrF* translation (16).

In addition, transcriptional control of *yscW-lcrF* has been predicted to be mediated by the <u>H</u>istone-like <u>N</u>ucleoid <u>s</u>tructuring protein, H-NS (16). H-NS contains an N-terminal oligomerization domain and a C-terminal DNA minor-groove binding domain separated by a flexible linker (18,19). H-NS preferentially binds AT rich regions of DNA (19,20). Once H-NS binds a high-affinity site, H-NS oligomerizes on the DNA (21,22). H-NS oligomers can either form a nucleoprotein filament on a contiguous stretch of DNA, or H-NS can form DNA bridges when multiple discrete H-NS binding regions are brought together, either way leading to transcriptional silencing of that particular gene (23). Interestingly, H-NS in multiple bacterial pathogens has been shown to silence certain gene targets during growth outside of the mammalian host (20-30°C), but fails to silence these same targets to the same magnitude when exposed to mammalian body temperature (37°C) (24–26). This suggests H-NS may play a role in repressing virulence factors outside host organisms in facultative pathogens. However, H-NS has been suggested to be an essential gene in pathogenic *Yersinia* (27,28), making it challenging to definitively test the role of H-NS in regulating gene expression in these organisms. Additionally, YmoA (*“<u>Y</u>ersinia* <u>mo</u>dul<u>a</u>tor”) in *Y. pseudotuberculosis*, an *E. coli* Hha (“<u>h</u>igh <u>h</u>emolysin <u>a</u>ctivity”) ortholog, has been suggested to modulate H-NS repression of a subset of promoters and deletion of *ymoA* in *Yersinia* leads to changes in gene expression of putative H-NS targets (16,29–31). YmoA and Hha lack a DNA binding domain; instead, these proteins form a heterocomplex with H-NS or H-NS paralogs (32–35). Recent data has suggested that Hha contributes to H-NS silencing by aiding in H-NS bridging (36). In the plague agent *Yersinia pestis*, YmoA is suggested to have a higher turnover rate at 37°C compared to environmental temperatures (30). While YmoA alone cannot bind the *yscW-lcrF* promoter, H-NS alone or the YmoA/H-NS complex can (16). Current models suggest that degradation of YmoA and therefore a reduction in the YmoA/H-NS complex at 37°C relieves repression of *yscW-lcrF* (30). Yet, *ymoA* deletion mutants exhibit even higher levels of T3SS expression at 37°C compared to a parental strain in all three pathogenic *Yersinia* species (16,29,30), suggesting that some YmoA is present even at 37°C during mammalian infection.

The <u>I</u>ron <u>S</u>ulfur <u>C</u>luster <u>R</u>egulator IscR is a critical positive regulator of *lcrF* (17,37). IscR belongs to the Rrf2 family of winged helix-turn-helix transcription factors (38,39). IscR was first characterized in *E. coli* where it exists in two forms: holo-IscR bound to a [2Fe-2S] cluster, and cluster-less apo-IscR (40–43). Both forms of IscR bind DNA, but while both apo-IscR and holo-IscR bind to so-called type II motif sequences, only holo-IscR binds type I motifs (41,42) Holo-IscR represses its own expression through binding two type I motifs in the *isc* promoter (44). Thus, conditions that increase iron-sulfur cluster demand, such as iron starvation or oxidative stress, lead to a lower holo- to apo-IscR ratio and higher overall IscR levels. *E. coli* IscR has been shown to activate or repress transcription of target genes *in vitro* and *in vivo* (41). We have previously shown that low iron and oxidative stress lead to upregulation of IscR in *Yersinia*, and subsequently upregulation of *lcrF* transcription and T3SS expression (17,37). Although we have shown IscR must bind upstream of the *yscW-lcrF* promoter to promote *lcrF* expression, the mechanism by which IscR promotes *lcrF* transcription is unknown. In this study, we find that IscR does not enhance *in vitro* transcription of *yscW-lcrF* mRNA. Instead, we show that IscR antagonizes YmoA/H-NS repression of the *yscW-lcrF* promoter to induce type III secretion.

## Results

### IscR does not promote *lcrF* transcription *in vitro*

IscR has previously been shown to enhance transcription by directly activating RNA polymerase activity or by antagonizing transcriptional repressors (41,45). To determine the molecular mechanism by which IscR potentiates transcription of *yscW-lcrF*, we performed an *in vitro* transcription assay with a DNA fragment containing the wild-type *Y. pseudotuberculosis yscW-lcrF* promoter (−143 to +58 bp relative to the +1 transcription start site) and the IscR binding site. Surprisingly, no change in *yscW-lcrF* transcription was observed after addition of apo-IscR (Fig 1). However, IscR was able to promote transcription of both *Y. pseudotuberculosis sufA* and *E. coli sufA*. These data suggest that IscR does not enhance *yscW-lcrF* transcription by regulating RNA polymerase directly. We therefore hypothesized that IscR promotes *yscW-lcrF* expression by antagonizing a repressor.

**Figure 1.**
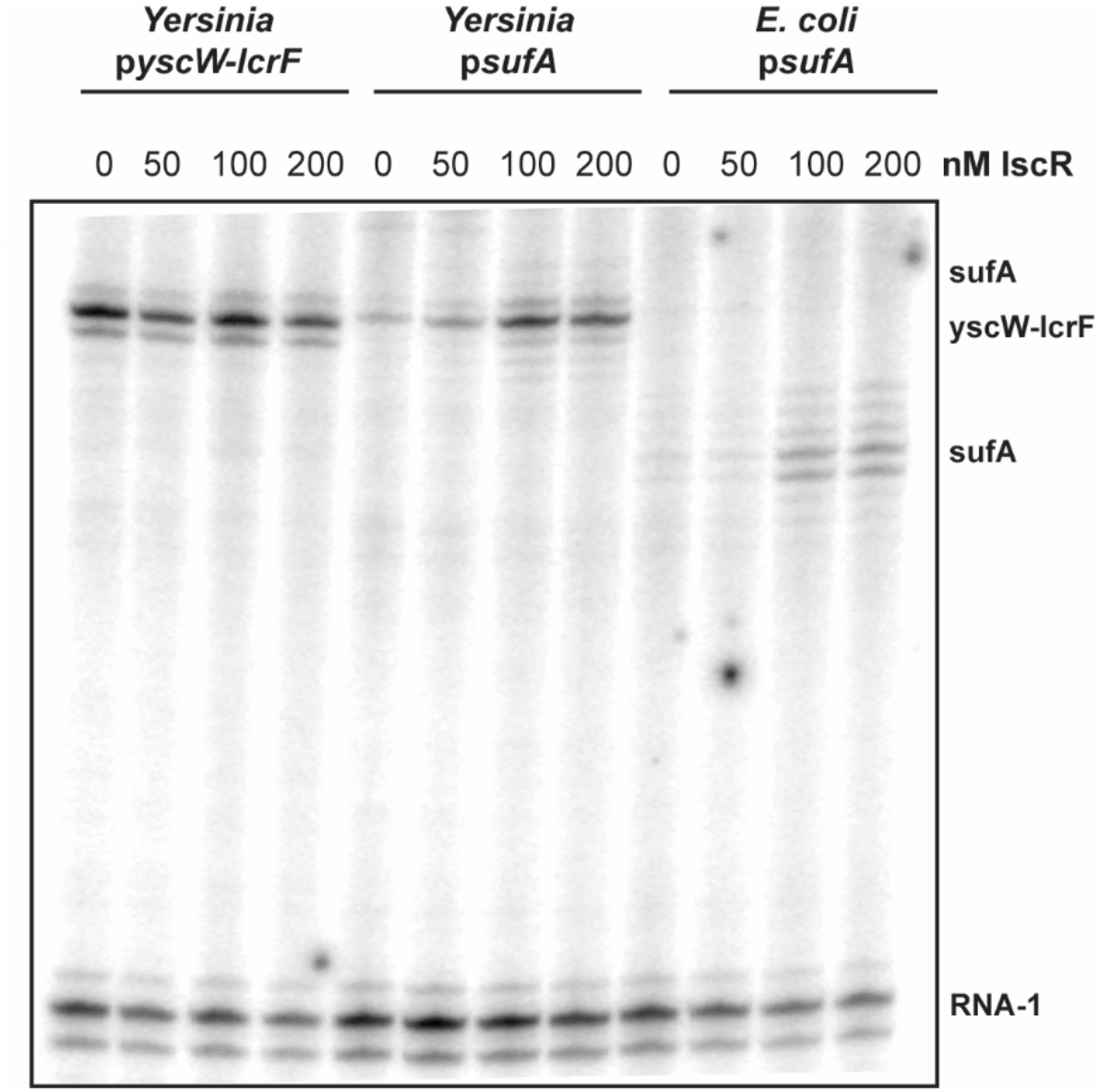
IscR does not directly promote transcription of *yscW-lcrF in vitro*. *In vitro* transcription reactions containing plasmids encoding the promoter of interest (*yscW-lcrF*, *sufA*, or *E. coli* K12 MG1655 *sufA*), Eσ70 RNA polymerase, and, where indicated, 50-200 nM IscR C92A protein lacking iron sulfur cluster coordination were incubated and analyzed. RNA-1 served as a control for this experiment.

### IscR is not required for LcrF expression or type III secretion in the absence of YmoA

Loss of *iscR* leads to a profound defect in T3SS activity while disruption of *ymoA* causes enhanced T3SS activity (16,17,37,46,47). We therefore hypothesized that IscR antagonizes YmoA-dependent repression of the T3SS. To test this, we assessed T3SS activity of *Y. pseudotuberculosis* expressing or lacking *iscR* and/or *ymoA*. Consistent with previous studies, we observed ~18-fold decrease in secretion of the T3SS effector protein YopE upon *iscR* deletion, while *ymoA* deletion led to ~6-fold increase in YopE secretion (Fig 2A). As expected for this transcriptional circuit, the effect of YmoA on T3SS activity required LcrF, the direct regulator of the T3SS (Fig S1). Importantly, YopE secretion in the Δ*iscR*/Δ*ymoA* double mutant was similar to Δ*ymoA* mutant, indicating that IscR is dispensable for T3SS activity in the absence of YmoA (Fig 2A).

**Figure 2.**
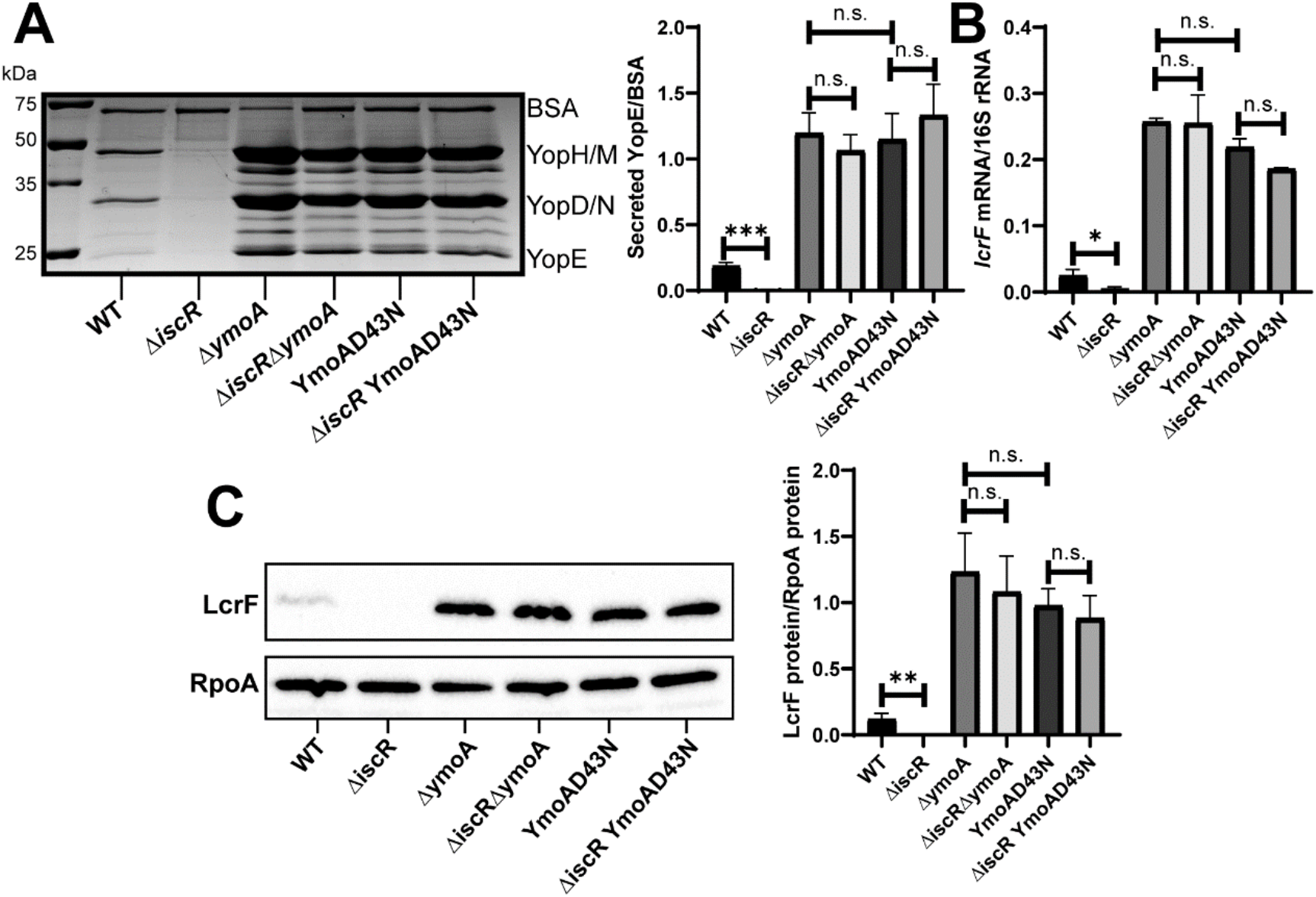
IscR is dispensable for type III secretion in the Δ*ymoA* mutant background. *Yersinia* strains were grown under T3SS-inducing conditions (low calcium at 37°C). (A) Precipitated secreted proteins were visualized by SDS-PAGE followed by Coomassie blue staining. Bovine serum albumin (BSA) was used as a loading control (left panel). Densitometry was used to measure the relative amount of secreted YopE T3SS effector protein versus BSA control. The average of four independent replicates ± standard deviation is shown (right panel). **(B)** RNA was extracted and reverse transcriptase quantitative PCR (RT-qPCR) was used to measure relative levels of *lcrF* mRNA normalized to 16S rRNA. The average of at least three biological replicates are shown ± standard deviation. **(C)** LcrF protein levels were determined by Western blotting (left panel) and densitometry (right panel) relative to the RpoA loading control. Shown is the average of four independent replicates ± standard deviation. Statistical analysis was performed using an unpaired Student’s t-test (*p<.05, **p<.01, ***p < .001, and n.s. non-significant).

Proteins of the YmoA family lack a DNA binding domain and are thought to affect transcription by its interaction with the histone like protein H-NS (31,36). Previous work has shown that a complex of YmoA/H-NS, but not YmoA alone, binds the *yscW-lcrF* promoter (16). To test the requirement for a YmoA/H-NS complex in regulation of YopE secretion by IscR we made use of a YmoA D43N mutant, which cannot interact with H-NS *in vitro* (48) and which we showed was produced in *Y. pseudotuberculosis* (Fig S2). Indeed, a *ymoA*^D43N^ mutant exhibited ~6-fold increase in YopE secretion similar to a *ymoA* deletion (Fig 2A). This suggests that YmoA represses *yscW-lcrF* through its interaction with H-NS. Furthermore, there was no difference in YopE secretion between the *ymoA^D43N^* mutant and the *iscR*/*ymoA^D43N^* double mutant. These effects on YopE secretion are most easily explained by changes in *lcrF* transcription and accordingly, LcrF protein levels. Indeed, while the Δ*iscR* mutant had a ~5-fold reduction in *lcrF* mRNA compared to wildtype and the Δ*ymoA* and *ymoA*^D43N^ mutants displayed ~10-fold elevated *lcrF* mRNA, we observed no difference in *lcrF* mRNA levels between the Δ*ymoA* and Δ*iscR*/Δ*ymoA* mutants (Fig 2B). Accordingly, we observed no difference in LcrF protein levels when *iscR* was deleted from the *ymoA* mutants (Fig 2C). Collectively, these data suggest that YmoA requires H-NS binding to inhibit *lcrF* transcription, and that IscR only exerts its positive effect on *lcrF* transcription in the presence of the YmoA/H-NS complex.

### IscR binding to the *yscW-lcrF* promoter is critical for LcrF expression only in the presence of YmoA

As IscR did not modulate YmoA or H-NS expression (Fig S3), we hypothesized that IscR must directly regulate LcrF by binding to the *yscW-lcrF* promoter to antagonize YmoA/H-NS-mediated repression. In order to test whether IscR binding to the *yscW-lcrF* promoter is important for regulating LcrF expression in the presence of YmoA, we used a previously characterized IscR binding site mutant (*lcrF*P^Null^) that ablates IscR binding to the *yscW-lcrF* promoter but expresses wildtype IscR (17). As expected, the *lcrF*^pNull^ exhibited a ~5-fold reduction in *lcrF* mRNA similar to what was observed in an *iscR* deletion mutant (Fig 3A). LcrF protein was completely undetectable in the *lcrF*^pNull^ compared to the wildtype strain (Fig 3B). However, in the absence of *ymoA*, this reduction in LcrF expression or T3SS activity by the *lcrF^pNull^* mutation was eliminated (Fig 3A-C). Taken together, these data suggest that IscR-dependent activation of LcrF expression in the presence of YmoA requires direct binding of IscR to the *yscW-lcrF* promoter.

**Figure 3.**
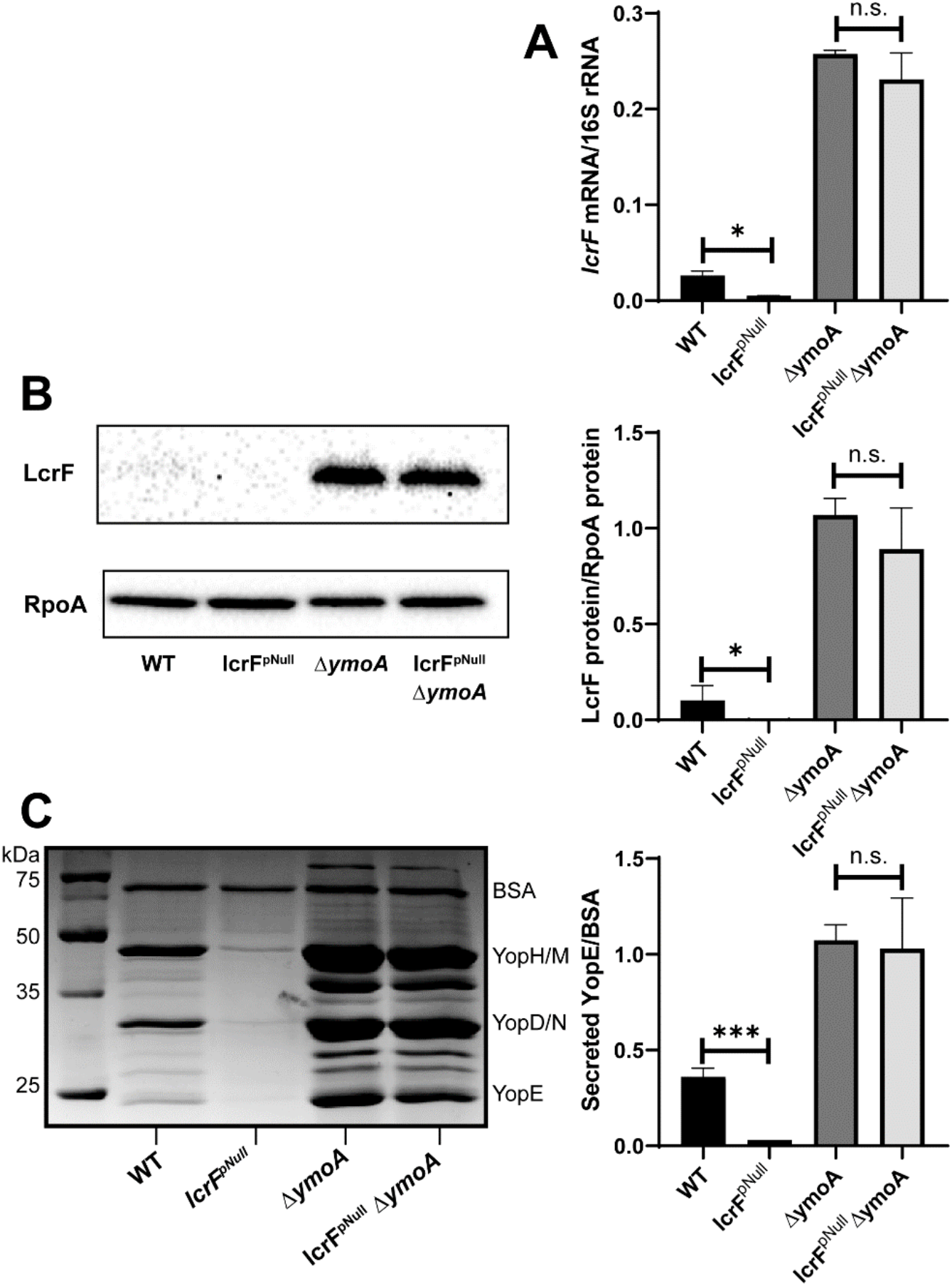
IscR binding to the *yscW-lcrF* promoter is dispensable in the absence of *ymoA*. *Yersinia* strains were grown under T3SS-inducing conditions. **(A)** Levels of *lcrF* mRNA were measured and normalized to 16S rRNA using RT-qPCR. The average of at least three biological replicates are shown ± standard deviation. **(B)** LcrF protein levels were measured relative to the RpoA loading control by Western blotting (left panel) and densitometry (right panel). Shown is the average of four biological replicates ± standard deviation. **(C)** Secreted proteins were precipitated and visualized by SDS-PAGE followed by Coomassie blue staining (left panel). The YopE bands were normalized to the BSA loading control (right panel). The average of three biological replicates ± standard deviations are shown. Statistical analysis was performed using an unpaired Student’s t-test (*p<.05, ***p < .001, and n.s. non-significant).

### Knockdown of H-NS leads to derepression of LcrF

We next examined the role of H-NS in the regulation of *lcrF* expression. H-NS has been proposed to be essential in both *Y. pseudotuberculosis* and *Y. enterocolitica* (27,28). Therefore, in order to test whether reducing H-NS occupancy at the *yscW-lcrF* promoter affects LcrF expression, we used CRISPRi to knockdown H-NS expression in wildtype *Y. pseudotuberculosis* and measured *lcrF* expression levels. For this CRISPRi system pioneered in *Yersinia pestis* (49), target gene guide RNAs and dCas9 can be induced in the presence of anhydrotetracycline (aTC). CRISPRi knockdown led to a ~6-fold decrease in H-NS transcription when exposed to aTC (Fig 4A). Importantly, this reduction of H-NS expression led to a ~31-fold increase in *lcrF* mRNA, suggesting H-NS represses LcrF transcription (Fig 4B). Knockdown of H-NS did not affect expression of *gyrA*, a housekeeping gene which is not predicted to be regulated by H-NS (Fig 4C). These data provide the first direct evidence that H-NS negatively influences *Yersinia* LcrF expression.

**Figure 4.**
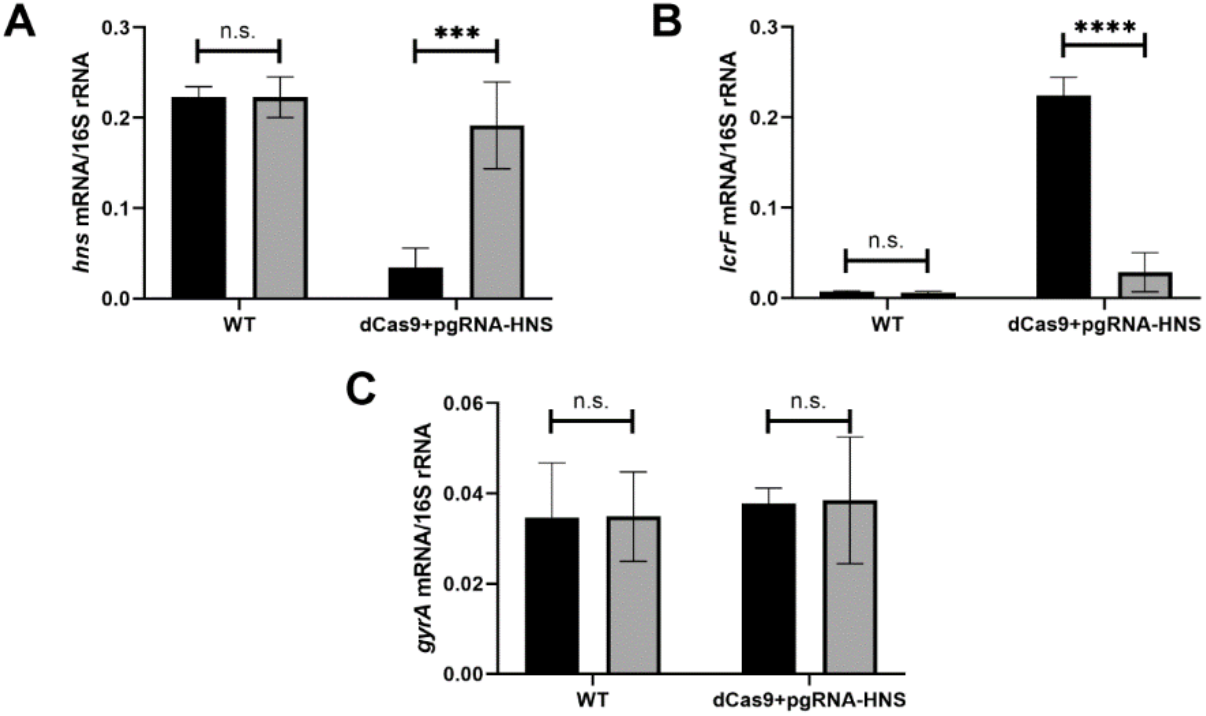
Knockdown of H-NS leads to derepression of LcrF. *Y. pseudotuberculosis* strains were grown in low calcium LB in the absence (grey bars) or presence (black bars) of 1μg/mL anhydrotetracycline for 3 hrs at 26°C to induce expression of *hns* guide RNA and dCas9 and then transferred to 37°C (T3SS inducing conditions) for 1.5 hrs. RNA was analyzed by RT-qPCR for *hns* **(A)**, *lcrF* **(B)**, or *gyrA* **(C)** mRNA level normalized to 16S rRNA. The average of three biological replicates are shown ± standard deviation. Statistical analysis was performed using an unpaired Student’s t-test (***p<.001, ****p<.0001, and n.s. non-significant).

### Two H-NS binding sites are required to repress *yscW-lcrF* promoter activity

H-NS and YmoA/H-NS complexes have been shown *in vitro* to bind the *yscW-lcrF* promoter between the −2 to the +272 position relative to the transcriptional start site (16). However, the exact H-NS binding site was not identified. We used FIMO-MEME suite tools to predict putative H-NS binding sites upstream of *yscW-lcrF* and identified three predicted H-NS binding sites (p-value<10^−3^; Fig 5A). These data suggested that H-NS may form a DNA bridge at this locus and repress *yscW-lcrF* transcription (50,51). To characterize which regions of the *yscW-lcrF* promoter allow for H-NS-YmoA repression and IscR activation, we systematically truncated the *yscW-lcrF* promoter and tested promoter activity using a *lacZ* reporter in the wildtype, Δ*iscR*, Δ*ymoA*, and Δ*iscR*/Δ*ymoA* backgrounds (Fig 5A). As expected, deletion of *ymoA* led to an increase in activity of the longest promoter construct, while *iscR* deletion led to a decrease in this promoter activity compared to the wildtype strain (Fig 5B). Consistent with our previous data showing that IscR was dispensable in the absence of YmoA, deletion of *iscR* in a Δ*ymoA* background did not inhibit the derepressed promoter activity seen in the Δ*ymoA* background. Eliminating the most upstream predicted H-NS binding site did not affect promoter activity (promoter 1 compared to promoter 2). However, additional truncation of the second H-NS binding site led to an increase in promoter activity in the wildtype and Δ*iscR* backgrounds, but not in the backgrounds lacking *ymoA* (promoter 2 compared to promoter 3) suggesting that some of the repressive effect by H-NS/YmoA had been lost. Importantly, further truncation to eliminate the IscR binding site led to deregulated promoter activity that was independent of IscR and YmoA (promoter 4). These data suggest that IscR is required to disrupt YmoA/H-NS repressive activity, explaining why it is dispensable in the absence of YmoA/H-NS. Lastly, truncation to eliminate the −35 and −10 promoter elements led to a complete lack of promoter activity (promoter 5). Taken together, these data suggest that IscR binding to the *ycsW-lcrF* promoter antagonizes YmoA/H-NS repression.

**Figure 5.**
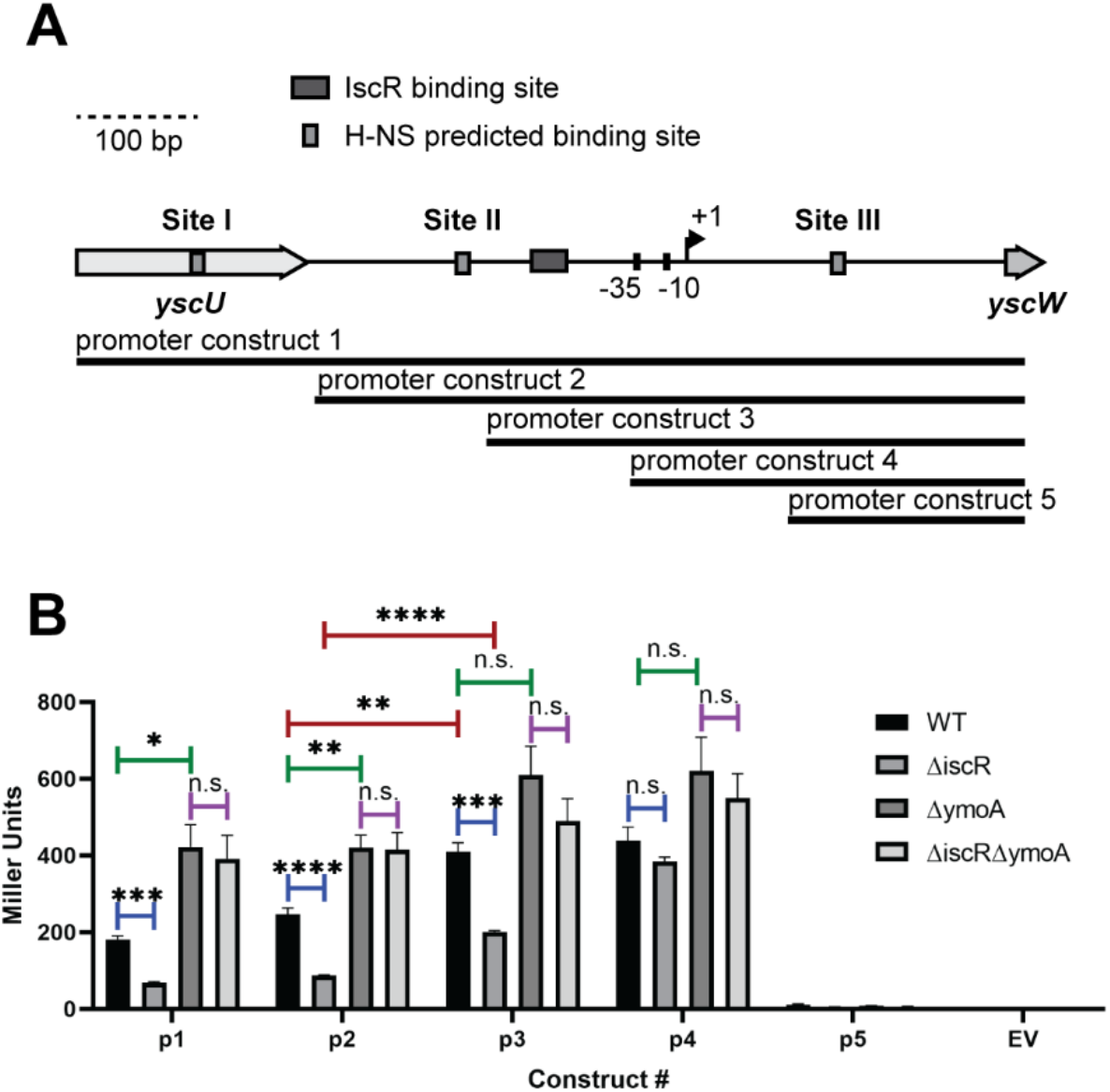
Defining regulatory regions required for IscR and H-NS-YmoA control of *PyscW-lcrF* expression. **(A)** Diagram of the *yscW-lcrF* promoter region (800 bp) containing the known IscR binding site (dark grey box), three MEME-suite FIMO predicted H-NS binding sites, referred to as Site I, Site II, and Site III (light grey boxes) and the previously characterized transcriptional start site (arrow). Schematic of P*yscW-lcrF::lacZ* fusions. Five constructs (p1-p5) were used to assess which regions of *pyscW-lcrF* allows for H-NS-YmoA repression and IscR activation. **(B)** *Yersinia* harboring the various *pyscW-lcrF::lacZ* plasmids were grown under T3SS-inducing conditions (low calcium LB at 37°C) for 1.5 hrs and assayed for β-galactosidase (Miller units). The average of at least three biological replicates are shown ± standard deviation. Statistical analysis was performed using an unpaired Student’s t-test (*p<.05, **p<.01, ***p<.001, ****p<.0001, and n.s. non-significant).

To test whether H-NS binds to these predicted sites, we carried out ChIP-qPCR analysis to assess H-NS occupancy at the I, II, and III putative binding regions *in vivo*. In order to immunoprecipitate H-NS-DNA complexes, we used a chromosomally-encoded 3xFLAG tagged H-NS allele. This FLAG tag did not affect the ability of H-NS to repress LcrF expression (Fig S4). Interestingly, previous reports have shown that H-NS in other facultative pathogens represses the expression of certain virulence genes under environmental temperatures (<30°C) but exhibits decreased binding at mammalian body temperature (37°C) (24,26,52). Consistent with this, we could not detect H-NS binding at any of these predicted sites at 37°C, but did observe H-NS enrichment at all three predicted sites in the *yscW-lcrF* promoter when bacteria were cultured at 26°C (Fig 6A). In contrast, no enrichment of H-NS was seen at a control pYV-encoded promoter that was not predicted to bind H-NS at either temperature (DN756_21750).

**Figure 6.**
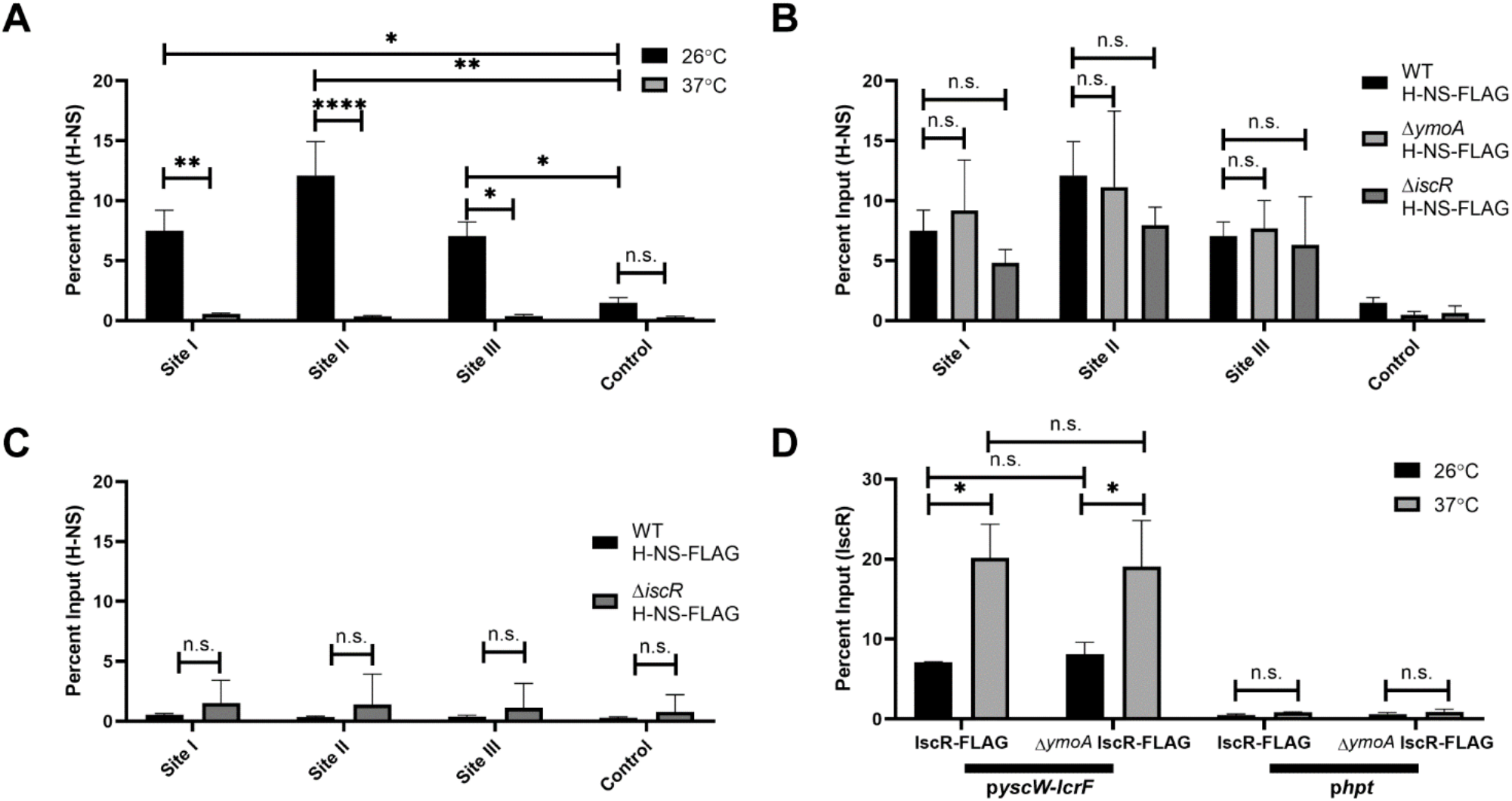
H-NS and IscR bind to the *yscW-lcrF* promoter at different temperatures. **(A)** The relative enrichment (percent input) of Site I, Site II, and Site III promoter DNA and a control promoter (DN756_21750), which H-NS is not predicted to bind, was analyzed by anti-FLAG ChIP-qPCR in *Yersinia* expressing H-NS-FLAG. ChIP-qPCR was performed with bacteria grown at 26°C (black bars) or 37°C (grey bars) in low calcium LB for 3 hrs. The average of at least three biological replicates ± standard deviation is shown. **(B)** ChIP-qPCR was performed with the H-NS-FLAG allele in wildtype, Δ*ymoA*, or Δ*iscR* mutant background at 26°C **(C)** ChIP-qPCR was performed with the H-NS-FLAG allele in wildtype or Δ*iscR* mutant background at 37°C. The average of at least three biological replicates ± standard deviation is shown. **(D)** ChIP-qPCR was performed with the IscR-FLAG allele in the wildtype or Δ*ymoA* mutant background at 26°C (black bars) or 37°C (grey bars). The *hpt* control promoter, which IscR is not predicted to bind, was used as a negative control. The average of at least three biological replicates ± standard deviation is shown and statistical analysis was performed using Two-way ANOVA (*p<.05, **p<.01, ***p<.001, ****p<.0001 and n.s. non-significant).

YmoA is predicted to affect the repressive ability of H-NS but was not shown to affect H-NS binding to the *yscW-lcrF* promoter (16). Consistent with this, no difference in H-NS binding was observed in the *ymoA* mutant compared to the parental strain at 26°C, suggesting that YmoA does not affect H-NS occupancy at the *yscW-lcrF* promoter at this temperature (Fig 6B). Likewise, there was no difference in H-NS enrichment at the *yscW-lcrF* promoter between the *iscR* mutant and the wildtype strain at 26 °C (Fig 6B) or 37 °C (Fig 6C). It is possible that H-NS binds to the *yscW-lcrF* promoter at 37°C, but this is below the limit of detection for ChIP-qPCR. Taken together, these data suggest that H-NS occupies the *yscW-lcrF* promoter at high levels under environmental temperatures at which the T3SS is repressed.

We also measured IscR enrichment at the *yscW-lcrF* promoter *in vivo*. We used a chromosomal 3xFLAG tagged IscR allele previously shown not to affect IscR activity (53). Interestingly, IscR enrichment at the *yscW-lcrF* promoter was ~3-fold higher at 37°C compared to 26°C (Fig 6D). This increase in IscR binding is not due to increased IscR levels since we do not observe higher levels of IscR protein when cultured at 37°C compared to 26°C (Fig S3), nor do we see increased binding of IscR at the promoter of another known IscR target, the *suf* operon (Fig S5). In addition, deletion of *ymoA* did not affect IscR occupancy at the *yscW-lcrF* promoter at 26°C or 37°C. These data suggest that at environmental temperatures, H-NS binds to the *yscW-lcrF* promoter at high levels and represses transcription, while at mammalian body temperature IscR binding to the *ycsW-lcrF* promoter antagonizes residual YmoA/H-NS-mediated repression.

### Environmental cues that increase IscR levels enable derepression of the *yscW-lcrF* promoter

We previously showed that low iron and high oxidative stress lead to elevated IscR levels, which then activate the T3SS through upregulation of LcrF (17). The data shown here suggest that this increase in IscR levels may be necessary to antagonize repressive YmoA-H-NS-activity at the *yscW-lcrF* promoter. To test this model, we measured *lcrF* mRNA levels in Δ*iscR* and Δ*ymoA* mutants under aerobic or anaerobic conditions at 37°C. As expected, under aerobic conditions *iscR* mRNA levels were increased ~4-fold compared to anaerobic conditions (Fig 7A). This upregulation of *iscR* levels led to a ~12-fold induction in *lcrF* levels in the wildtype strain (Fig 7B). In contrast, *lcrF* mRNA and protein levels were not affected by oxygen in the Δ*ymoA* and Δ*iscR*/Δ*ymoA* mutants (Fig 7A, 7C). It is important to note that deletion of *ymoA* reduced expression of IscR mRNA and protein expression under these conditions, although this does not explain elevated LcrF/T3SS expression in the *ymoA* mutant. Indeed, IscR occupancy at the *lcrF* promoter is not affected by *ymoA* deletion (Fig 6D). Taken together, these data suggest that environmental conditions that increase IscR levels (such as aerobic conditions) disrupt YmoA/H-NS-mediated repression of *lcrF* expression.

**Figure 7.**
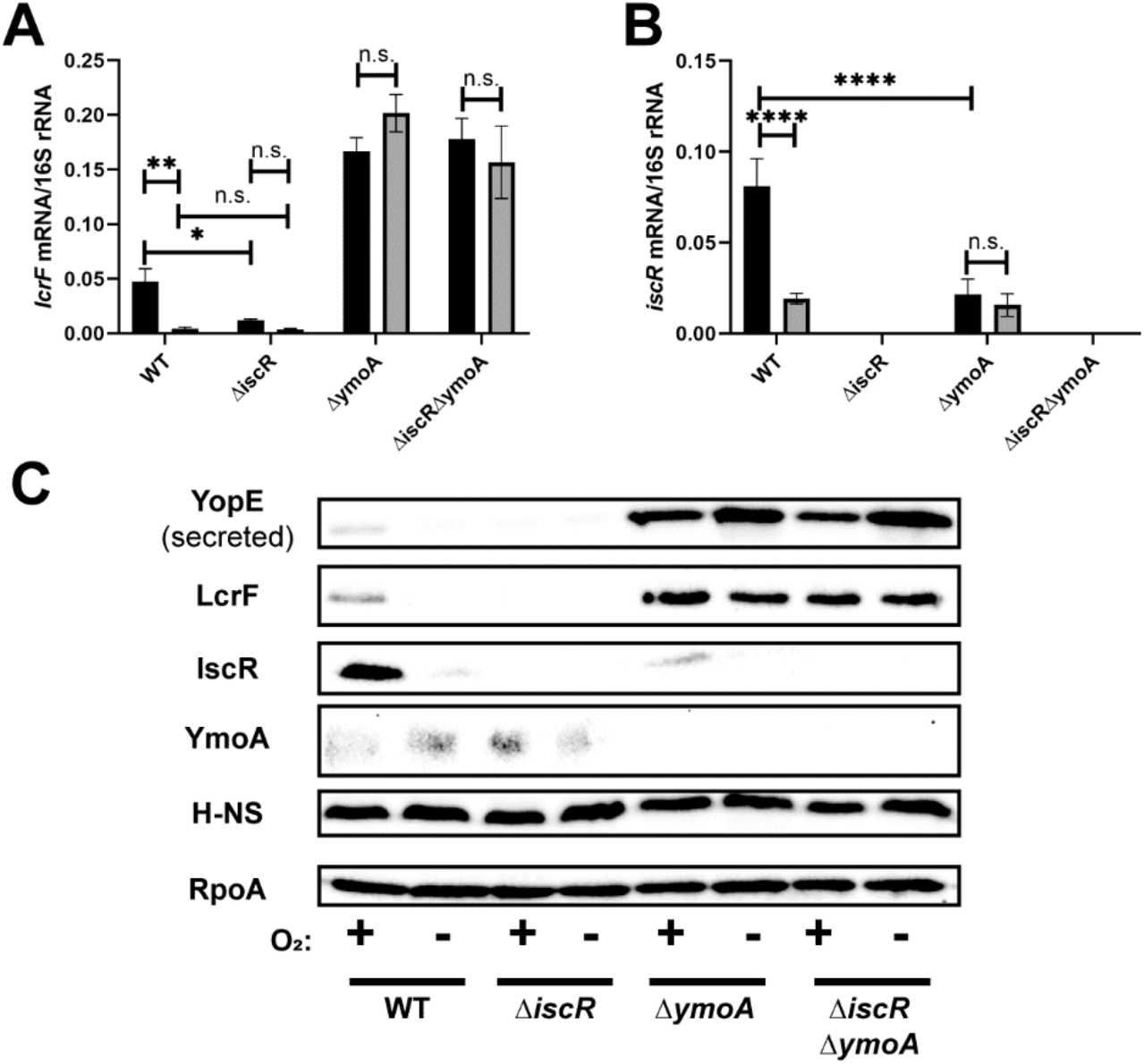
Oxygen-dependent control of *lcrF* requires YmoA. *Yersinia* strains were cultured under T3SS inducing conditions under aerobic (black bars) or anaerobic (grey bars) conditions. Levels of *lcrF* **(A)** and *iscR* **(B)** mRNA levels were measured by RT-qPCR and normalized to 16S rRNA. The average of at three biological replicates are shown ± standard deviation. **(C)** *Yersinia* strains were grown under similar conditions as stated above and whole cell extracts were probed for RpoA, IscR, H-NS, LcrF, YopE, and YmoA by Western blotting. One representative experiment out of three biological replicates is shown. Statistical analysis was performed using a one-way ANOVA with Tukey multiple comparisons (*p<.05,**p<.01,****p<.0001, and n.s. non-significant).

## Discussion

Our data suggests IscR activates transcription of *yscW-lcrF* by antagonizing repressive activity of YmoA-H-NS (Fig 8). Knockdown of *hns* expression by CRISPRi revealed that H-NS, a putative essential gene in *Yersinia*, is required for repression of *yscW-lcrF*. Furthermore, YmoA must interact with H-NS to repress *yscW-lcrF* transcription and overall T3SS activity at 37°C. Importantly, IscR promotes *yscW-lcrF* expression and T3SS activity only in the presence of YmoA-H-NS repression. Our data point to a model where H-NS occupies the *yscW-lcrF* promoter at environmental temperatures independently of YmoA and IscR, but at mammalian body temperature YmoA binding to H-NS represses the *yscW-lcrF* promoter only when IscR levels are low (Fig 8). *Y. pseudotuberculosis* IscR levels are thought to be kept low in the intestinal lumen, under anaerobic iron-replete conditions. Under these conditions, where the T3SS is not required for colonization, YmoA and H-NS cooperate to repress LcrF expression. Once *Yersinia* cross the intestinal barrier, oxygen tension increases and iron is scarce, allowing elevated IscR levels that antagonize YmoA/H-NS activity to allow LcrF expression and type III secretion, which is required for extraintestinal infection (54–57). This suggests that YmoA/H-NS and IscR work together to allow temperature and oxygen tension/iron availability to limit T3SS activity not just to only inside the host organism, but to only in extraintestinal tissue. Given that IscR is essential for T3SS activity in the related plague agent *Y. pestis* that does not enter the intestinal tract (17), we predict that in the flea vector that maintains temperatures lower than the mammalian host, H-NS represses LcrF expression. Then upon entry into the mammalian host bloodstream, the elevated temperature leads to decreased occupancy of YmoA/H-NS at the *yscW-lcrF* promoter to levels that allow IscR to antagonize YmoA/H-NS repression and facilitate expression of the T3SS activity required for early stages of plague (58,59).

**Figure 8.**
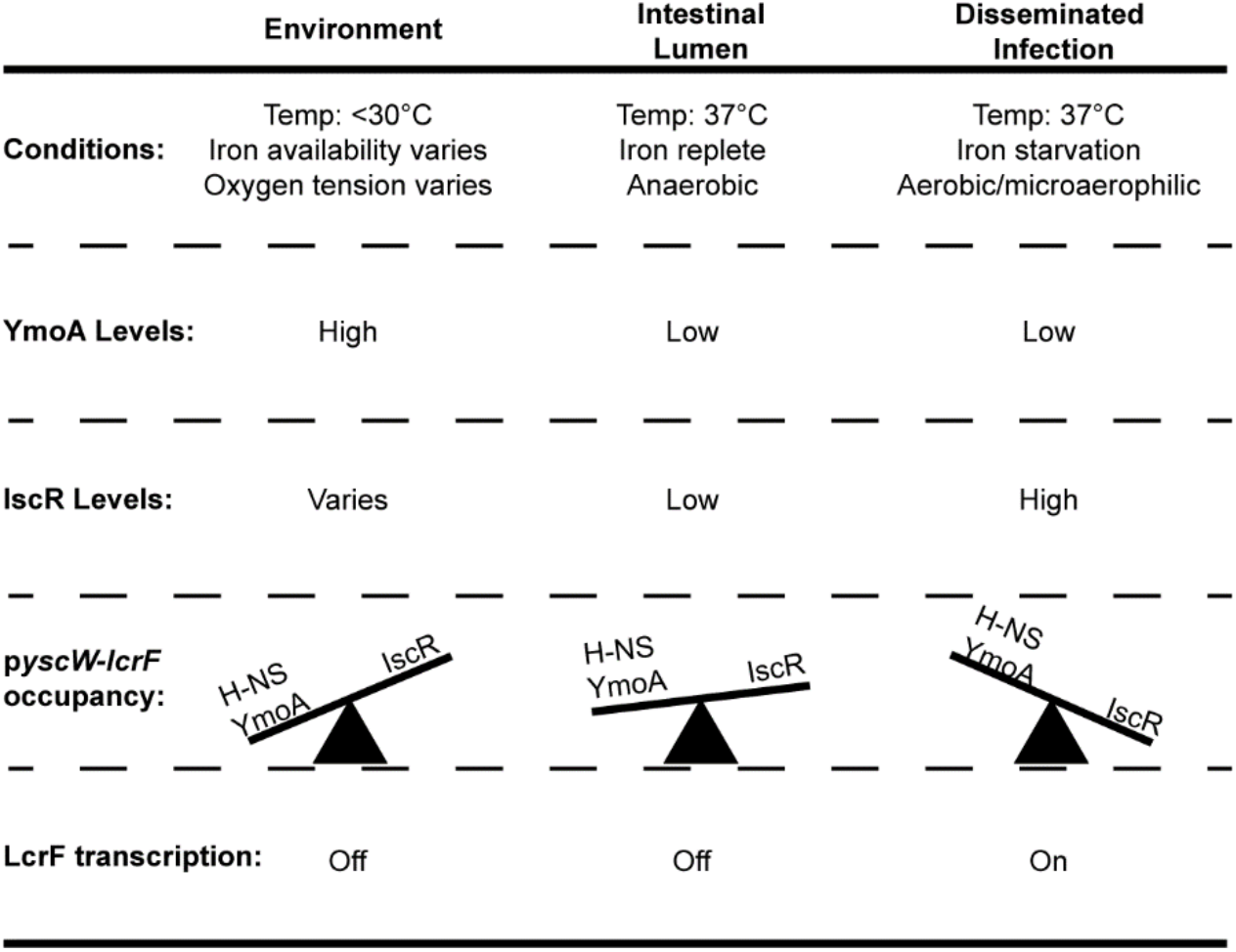
Proposed model for activation of *yscW-lcrF* via IscR. At environmental conditions (26°C, low oxidative stress, high iron), H-NS occupancy at the *yscW-lcrF* promoter is high and LcrF expression and T3SS activity is repressed. When *Y. pseudotuberculosis* becomes ingested, it travels to the intestinal lumen. YmoA protein levels decrease due to ClpXP/Lon protease activity at 37°C, but sufficient levels remain to potentiate H-NS-mediated repression of *yscW-lcrF* and prevent type III secretion. This is because IscR levels are kept low by anaerobic and iron replete conditions. Once *Y. pseudotuberculosis* crosses the intestinal barrier, it encounters higher oxygen tension and lower iron availability, causing an increase in IscR protein levels that can antagonize YmoA/H-NS repression of *yscW-lcrF* and allow for LcrF expression and T3SS activity.

Previous reports have suggested that H-NS targets a subset of genes under environmental conditions, but no longer represses those same genes under mammalian body temperature. For example, the *Shigella flexneri* T3SS is regulated by an AraC transcriptional regulator called VirF and H-NS (60,61). VirF promotes VirB, which ultimately activates the *Shigella* T3SS (62). The *Shigella* T3SS is only expressed under mammalian body temperature and this is controlled by preventing expression of VirF at environmental temperatures. Interestingly, H-NS was shown to directly repress *virF* transcription by binding to the *virF* promoter (52). Later studies found that H-NS binds to two distinct sites upstream of *virF* leading to the formation of a DNA bridge (63). This study also found that H-NS binds to a higher degree at the *virF* promoter under lower temperatures (<30°C) compared to mammalian body temperature (37°C). Thus H-NS was shown to repress promoter activity of *virF* at both lower temperatures (<30°C) and mammalian body temperature (37°C), however H-NS has a stronger effect on repression of *virF* under lower temperatures (52). This molecular mechanism is similar to what we report here, where H-NS occupies the *yscW-lcrF* promoter at environmental conditions and is below the limit of detection by ChIP-qPCR at 37°C. However, *Yersinia* H-NS repression of LcrF still occurs at 37°C unless IscR levels increase sufficiently to antagonize this repression.

YmoA was previously shown to bind to H-NS and the YmoA/H-NS complex was proposed to regulate LcrF expression (16,48). Indeed, a YmoA mutation that eliminates H-NS binding phenocopied the *ymoA* deficient strain, suggesting YmoA must interact with H-NS to repress LcrF/T3SS. However, YmoA was not shown to affect H-NS binding to the *yscW-lcrF* promoter *in vitro* (16), and our ChIP-qPCR analysis did not find a change in H-NS *yscW-lcrF* promoter occupancy in the presence or absence of YmoA at 26°C. At 37°C, H-NS occupancy was below the limit of detection by ChIP-qPCR, so we could not rule out H-NS or YmoA/H-NS binding to the *yscW-lcrF* promoter at this temperature. However, deletion of *ymoA* or knockdown of *hns* both caused elevated LcrF expression at 37°C, indicating that both proteins are needed to repress the *yscW-lcrF* promoter at mammalian body temperature. Although YmoA and the YmoA homolog Hha have been shown to bind DNA *in vitro*, YmoA and Hha lack a DNA binding domain and most likely purification of YmoA or Hha leads to copurification of H-NS or H-NS paralogs. This may explain why YmoA has been shown to interact with specific segments of DNA *in vitro*. Taken together, these data suggest that YmoA binding to H-NS does not alter H-NS occupancy at the *lcrF* promoter but, rather, potentiates H-NS repressive activity. *E. coli* Hha influences H-NS bridging and promotes H-NS silencing of target genes (36). We hypothesize that H-NS bound to YmoA forms a bridging complex on the *lcrF* promoter that represses *lcrF* transcription, and IscR disrupts this repressive complex. Testing this hypothesis will be the subject of future work.

The mechanism by which IscR promotes or represses transcription of target genes varies. For example, elevated transcription of *ydiU*, a gene of unknown function, in *E. coli* is thought to be driven by direct interaction between RNA polymerase and IscR (41). However, IscR has also been shown to activate transcription of other target genes by antagonizing a repressor (45). For example, in *Vibrio vulnificus* IscR promotes expression of the *vvhBA* operon, which encodes an extracellular pore-forming toxin essential for its hemolytic activity (45,64,65), while the *vvhBA* operon is repressed by H-NS (66). In *V. vulnificus*, nitrosative stress and iron starvation lead to upregulation of IscR (45). This increase in IscR leads to upregulation of *vvhBA* by increasing IscR levels and antagonizing H-NS repression of *vvhBA*. This molecular mechanism is very similar to what we observe here for IscR and H-NS in *Yersinia*, where aerobic conditions promote high IscR levels that antagonize H-NS repressive activity at the *yscW-lcrF* promoter. Therefore, antagonizing H-NS repression may be a common mechanism of gene regulation by IscR in response to changes in iron and oxygen.

## Materials and Methods

### Bacterial strains and growth conditions

Bacterial strains used in this paper are listed in Table S1 *Y. pseudotuberculosis* were grown, unless otherwise specified, in LB (Luria Broth) at 26°C shaking overnight. To induce the T3SS, overnight cultures were diluted into low calcium LB medium (LB plus 20 mM sodium oxalate and 20 mM MgCl_2_) to an optical density (OD_600_) of 0.2 and grown for 1.5 h at 26°C shaking followed by 1.5 h at 37°C to induce Yop synthesis, depending on the assay, as previously described (1).

For growing *Yersinia* under varying oxygen conditions, casamino acid-supplemented M9 media, referred to as M9 below, was used (2). Growth of cultures to vary oxygen tension was achieved by first diluting 26°C overnight aerobic cultures of *Y. pseudotuberculosis* to an OD_600_ of 0.1 in fresh M9 minimal media supplemented with 0.9% glucose to maximize growth rate and energy production under anaerobic conditions, and incubating for 12 hrs under either aerobic or anaerobic conditions at 26°C. Both aerobic and anaerobic cultures were diluted to an OD_600_ of 0.1, grown for 2 hrs at 26°C, and then shifted to 37°C for 4 hrs.

### Construction of *Yersinia* mutant strains

The *Yersinia* mutants were generated as described in (3). H-NS was tagged with a C-terminal 3xFLAG affinity tag at the native locus through splicing by overlap extension (4), using primer pair F*hns*_cds/R*hns*_cds (Table S2) to amplify ~500bp upstream of *hns* plus the *hns* coding region excluding the stop codon, F3×FLAG/R3×FLAG to amplify the 3×FLAG tag, and F3’*hns*/R3’*hns* to amplify the ~500 bp downstream region of *hns* including the stop codon. For the Δ*ymoA* mutant, primer pairs F5/R5Δ*ymoA* were used to amplify ~1000 bp 5’ of *ymoA* and F3/R3Δ*ymoA* to amplify ~1000 bp 3’ of *ymoA*. To generate the *ymoA*^D43N^ mutant, primer pairs pUC19_YmoA_F and pUC19_YmoA_R were used to amplify 250 bp upstream of *ymoA* to 250 downstream of the *ymoA* start codon and the amplified product cloned into a BamHI and SacI digested pUC19 plasmid. Q5 site directed mutagenesis was performed using primer pairs *ymoA*^D43N^ _F and *ymoA*^D43N^ _R. The resulting plasmid, pUC19 *ymoA*^D43N^, was digested with BamHI and SacI and the resulting fragment was ligated into the suicide plasmid pSR47s. Mutant strains were generated as described above.

In order to generate *lacZ* promoter constructs of *ymoBA* and *hns*, primer pairs pFU99a_ymoA_F/pFU99a_ymoA_R and pFU99a_hns_F/pFU99a_hns_R were used to amplify ~500 bp upstream of *ymoA* and *hns*, respectively, which included the first ten amino acids of *ymoA* and *hns*. These promoters and first ten amino acids of YmoA and H-NS were fused in frame to *lacZ* and cloned into a BamHI- and SalI-digested pFU99a using the NEBuilder HiFi DNA Assembly kit (New England Biolabs, Inc) electroporated into *Y. pseudotuberculosis*.

In order to generate *lacZ* promoter constructs of *yscW-lcrF*, the reverse primer pFU99a_*yscWlcrF*_R was used with the following forward primers: pFU99a_*yscWlcrF*_p1 (promoter construct 1/−505 to +294 of *yscW*), pFU99a_*yscWlcrF*_p2 (promoter construct 2/ −309 to +294 of *yscW*), pFU99a_*yscWlcrF*_p3 (promoter construct 3/ −166 to +294 of *yscW*), pFU99a_*yscWlcrF*_p4 (promoter construct 4/ −47 to +294 of *yscW*), or pFU99a_*yscWlcrF*_p5 (promoter construct 5/ +101 to +294 of *yscW*). These promoter fragments were cloned into a BamHI- and SalI-digested pFU99a and electroporated into *Y. pseudotuberculosis*.

### *In vitro* transcription assay

The DNA template used to assess if IscR could directly promote transcription of the *yscW-lcrF* promoter contained the −147 to +53 bp relative to the +1 transcription start site of *yscW*. The promoter containing the *Y. pseudotuberculosis sufA* promoter and the *E. coli sufA* promoter, which IscR has been shown to directly promote transcription of, served as a positive control (3, 5). The effect of IscR-C92A on σ70-dependent activity was determined by incubating IscR-C92A with 2 nM supercoiled pPK12778 [purified with the QIAfilter Maxi kit (Qiagen)], 0.25 μCi of [α-32P]UTP (3,000 μCi/mmol; Perkin Elmer), 20 μM UTP, and 500 μM each of ATP, GTP, and CTP for 30 min at 37°C in 40 mM Tris (pH 7.9), 30 mM KCl, 10 mM MgCl2, 100 μg/mL bovine serum albumin (BSA), and 1 mM DTT. Purified apo-IscR was used since the IscR binding site upstream of *yscW-lcrF* has been characterized to be an IscR type II site, which apo-IscR is capable of binding (6, 7). Eσ70 RNA polymerase (NEB) was added to a final concentration of 50 nM and the reaction was terminated after 5 min by addition of Stop Solution (USB Scientific). Samples were heated for 60 s at 90°C, and loaded onto a 7 M urea-8% polyacrylamide gel in 0.5× Tris-borate-EDTA (TBE) buffer. The reaction products were visualized by phosphorimaging.

### Type III secretion system secretion assay

Visualization of T3SS cargo secreted in broth culture was performed as previously described (8). Briefly, *Y. pseudotuberculosis* in LB low calcium media (LB plus 20 mM sodium oxalate and 20 mM MgCl2) was grown for 1.5 h at 26°C followed by growth at 37°C for 1.5 h. Cultures were normalized to OD_600_ and pelleted at 13,200 rpm for 10 min at room temperature. Supernatants were removed and proteins precipitated by addition of trichloroacetic acid (TCA) at a final concentration of 10%. Samples were incubated on ice for at least 1 hr and pelleted at 13,200 rpm for 15 min at 4°C. Resulting pellets were washed twice with ice-cold 100% acetone and resuspended in final sample buffer (FSB) containing 0.2 M dithiothreitol (DTT). Samples were boiled for 5 min prior to separating on a 12.5% SDS-PAGE gel. Coomassie stained gels were imaged using Bio-Rad Image Lab Software Quantity and Analysis tools. YopE bands were quantified using this software and normalized to the BSA protein precipitation control.

### Western Blot Analysis

Cell pellets were collected, resuspended in FSB plus 0.2 M DTT, and boiled for fifteen minutes. At the time of loading, supernatants and cell pellets were normalized to the same number of cells. After separation on a 12.5% SDS-PAGE gel, proteins were transferred onto a blotting membrane (Immobilon-P) with a wet mini trans-blot cell (Bio-Rad). Blots were blocked for an hour in Tris-buffered saline with Tween 20 and 5% skim milk, and probed with the rabbit anti-RpoA (gift from Melanie Marketon), rabbit anti-LcrF (gift from Gregory Plano), rabbit anti-IscR (9), rabbit anti-YmoA (gift from Gregory Plano), rabbit anti H-NS (gift from Robert Landick), mouse M2 anti-FLAG (Sigma), goat anti-YopE (Santa Cruz Biotech), and horseradish peroxidase-conjugated secondary antibodies (Santa Cruz Biotech). Following visualization, quantification of the bands was performed with Image Lab software (Bio-Rad).

### Quantitative RT-PCR

RT-qPCR was carried out as previously described (3) using the primers in Table S2. The expression levels of each target gene were normalized to that of 16S rRNA present in each sample and calculated by utilization of a standard curve. At least three independent biological replicates were analyzed for each condition.

### β-galactosidase Assays

*Y. pseudotuberculosis* harboring promoter-*lacZ* fusion plasmids were grown in LB low calcium media (LB plus 20 mM sodium oxalate and 20 mM MgCl2) for 1.5 h at 26°C followed by growth at 37°C for 1.5 h. Protein expression was stopped by incubating cells on ice for 20 minutes. Cultures were spun down and resuspended in Z Buffer (10). Samples were permeabilized using chloroform and 0.1% sodium dodecyl sulfate, incubated with 0.8 mg/mL ONPG, and β-galactosidase enzymatic activity was terminated by the addition of 1M sodium bicarbonate. β-galactosidase activity is reported as Miller units.

### CRISPRi knockdown

Knockdown of H-NS via CRISPRi methods was adapted from (11). In order to generate the pgRNA-tetO-JTetR-H-NS plasmid, a protospacer-adjacent motif (PAM) was located near the promoter of *hns* (12). Two oligonucleotides (*hns*_gRNA_F and *hns*_gRNA_R) consisting of 20-nt targeting the *hns* promoter region with BbsI cohesive ends were synthesized and annealed before being cloned into pgRNA-*tetO*-JTetR by Golden Gate assembly. The plasmids pdCas9-bacteria and pgRNA-tetO-JTetR-H-NS were transformed into WT *Y. pseudotuberculosis* sequentially. These plasmids induce expression of dCas9 and gRNA-H-NS when exposed to anhydrotetracycline. *Y. pseudotuberculosis* cultures carrying these plasmids were sub-cultured to OD_600_ 0.2 and incubated at 26°C for 3 hrs in the presence or absence of 1μg/mL anhydrotetracycline, and then transferred to 37°C for 1.5 hrs to induce the T3SS. Samples were collected, and RNA was isolated for qRT-PCR analysis.

### Bioinformatic prediction of YmoA/H-NS binding sites

A training set of known H-NS binding sites in *E. coli* K-12 substr. MG1655 from RegulonDB was used to generate an H-NS binding motif using MEME-suite 5.1.1 tools (13, 14). FIMO was then used to scan for an H-NS binding site near the regulatory region of the *yscW-lcrF* promoter.

### ChIP-qPCR

Cells were grown for 3hrs at 26°C or 37°C with shaking at 250 rpm and protein/nucleic acids were crosslinked using 1% formaldehyde at 26°C or 37°C for 10 min. Crosslinking was quenched with the addition of ice cold 0.1 M glycine and incubated at 4°C for 30 min. 32×1OD_600_ cells were harvested for each replicate and cell pellets were stored at −80°C. DNA was fragmented by resuspending samples using IP buffer (100mM Tris-HCl, pH 8, 300mM NaCl, 1% Triton X-100, 1 mM PMSF) and sonicated at 25% Amplitude 15s on/ 59s off for a total of 8 cycles per sample. After sonication, lysates were treated with micrococcal nuclease and RNase-A for 1hr at 4°C. Lysates were clarified via centrifugation at 13,000 rpm for 15 min at 4°C. Lysates were pre-cleared using Dynabeads Protein A/G for 3hr at 4°C. Immunoprecipitation was performed by adding Sigma monoclonal mouse anti-FLAG M2 antibody to samples and incubated overnight at 4°C. Dynabeads Protein A/G were added to samples and washes were performed to remove non-specific binding. After H-NS-DNA or IscR-DNA complexes were eluted, samples were placed at 65°C for 5 hr to reverse crosslinks. DNA was then purified using Qiagen PCR purification kit and input samples were diluted 1:100 while samples treated with antibody or control samples not treated with the antibody were diluted 1:5 and qPCR was performed to assess IscR/H-NS binding to promoters of interest. Percent input was calculated by the following equation: 100*2^CT_input_ - CT^_+AB_.

## Data Availability

All study data are included in the article and SI Appendix. All experimental data will be made available upon request.

## Acknowledgments

We thank Gregory V. Plano (University of Miami Health System) for the YmoA antibody and Robert Landick (University of Wisconsin, Madison) for the H-NS antibody.

## Supporting Information

Figure S1. YmoA affects LcrF dependent type III secretion activity.

Figure S2. YmoA mutations do not affect mRNA levels or protein levels of IscR or H-NS.

Figure S3. IscR does not regulate YmoA or H-NS expression.

Figure S4. 3xFLAG tag allows for detection of H-NS using FLAG antibody and does not affect H-NS ability to repress LcrF

Figure S5. IscR enrichment at the *suf* promoter is not influenced by temperature.

Table S1. Strains used in this study.

Table S2. *Y. pseudotuberculosis* primers used in this study.

Table S3. Plasmids used in this study.

