## Supplemental material for "The transcription factor IscR promotes *Yersinia* type III secretion system activity by antagonizing the repressive H-NS-YmoA histone-like protein complex"

**This PDF file includes:**

Figures S1 to S5
Table S1 to S3
SI References

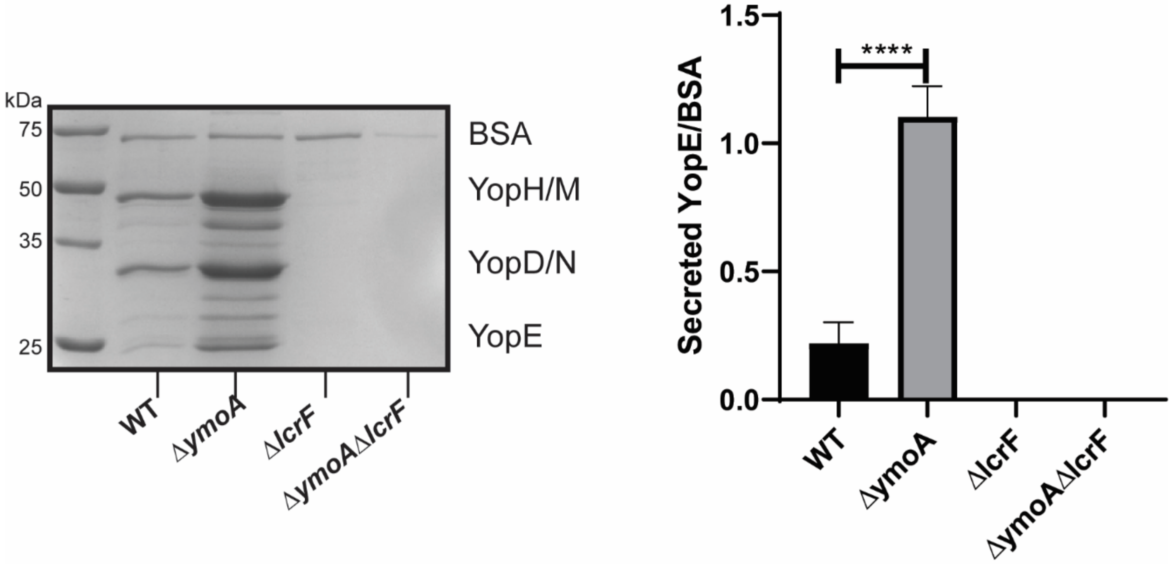


**Figure S1. YmoA affects LcrF dependent type III secretion activity.** *Yersinia* strains were grown in low calcium LB for 1.5 hrs at 26°C and transferred to 37°C (T3SS inducing conditions) for 1.5 hrs. The supernatant was collected and separated on a 12.5% SDS polyacrylamide gel, and subsequently stained with Coomassie blue (left panel). Bovine serum albumin (BSA) was used as a loading control. Gel bands were quantified by using Bio-Rad Image Lab Software Quantity and Analysis tools. YopE bands were normalized to the BSA loading control (right panel). The average of three independent replicates ± standard deviation is shown and statistical analysis was performed using an unpaired Student's t-test. (****p<.0001).

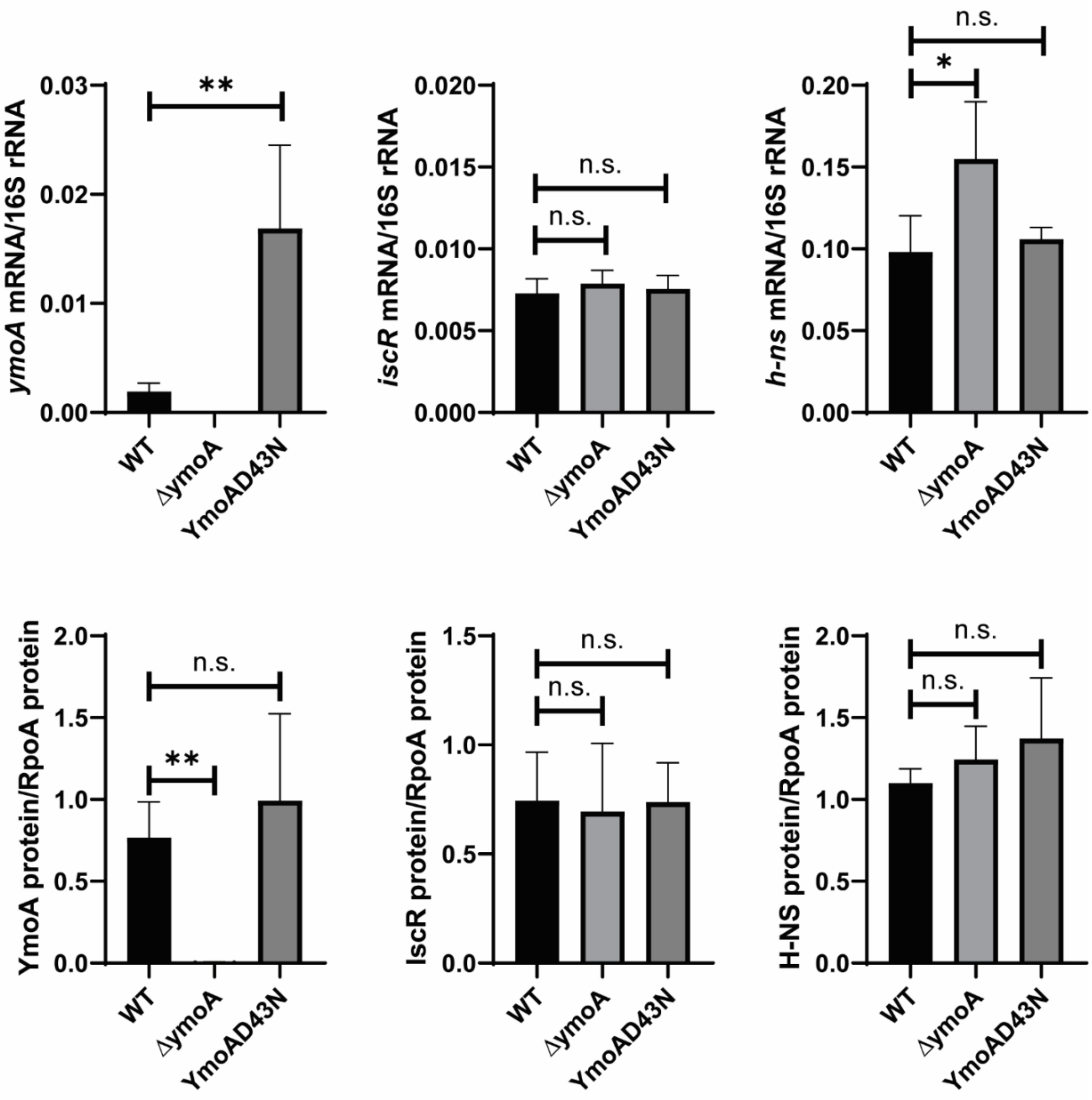


**Figure S2. YmoA mutations do not affect mRNA levels or protein levels of IscR or H-NS***.* RNA and whole cell extracts were prepared from indicated *Yersinia* strains grown in low calcium LB for 1.5 hrs at 26°C and transferred to 37°C (T3SS inducing conditions) for 1.5 hrs. *ymoA, iscR, and h-ns* mRNA expression levels was measured by RT-qPCR and normalized to 16S rRNA. The average of at least three biological replicates are shown with standard deviation (top panel). For western blots of cell extracts, proteins were visualized using anti-YmoA, anti-IscR, and anti-H-NS antibodies. Proteins were quantified by densitometry using BioRad Image Lab. The average of three biological replicates are shown with standard deviation (bottom panel). Statistical analysis was performed using an unpaired Student's t-test. (*p<.05, **p<.01, and n.s. non-significant).

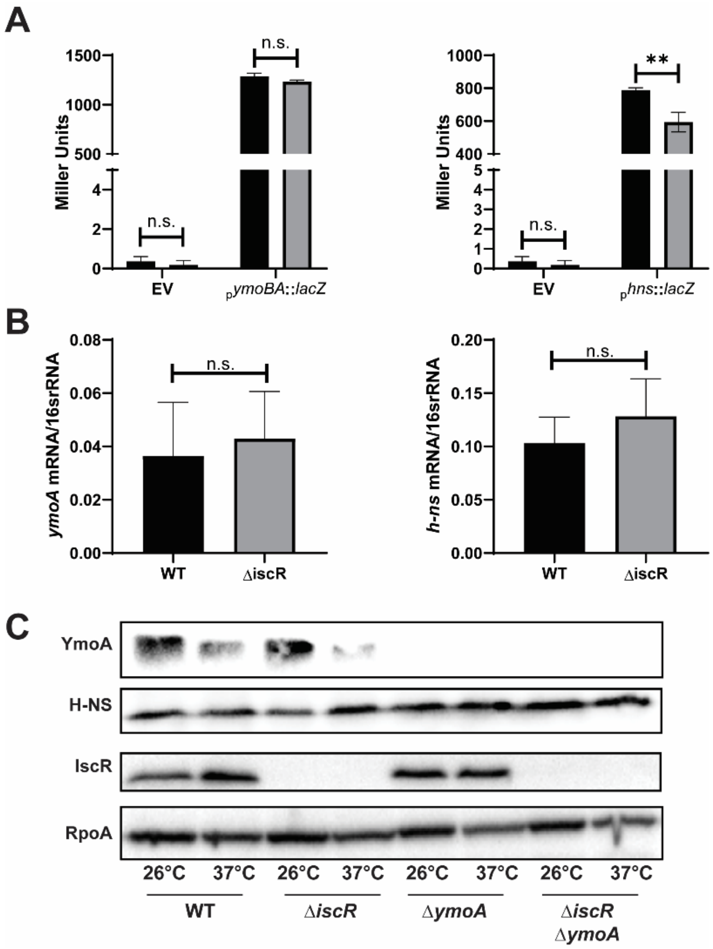


**Figure S3. IscR does not regulate YmoA or H-NS expression. (A)** *Yersinia* strains harboring either the vector pFU99a (EV) or pFU99a plasmid encoding the *ymoBA* or *hns* promoters fused to *lacZ* were grown under T3SS inducing conditions and assayed for β-galactosidase activity (Miller units). Black bars represent WT background and grey bars represent the *iscR* mutant background. The average of three biological replicates are shown ± standard deviation. (**B)** RNA was extracted from *Yersinia* strains grown under T3SS-inducing conditions and RT-qPCR was used to measure relative *ymoA* and *hns* mRNA levels normalized to 16S rRNA. Black bars represent WT background and grey bars represent the *iscR* mutant background. The average of at least three biological replicates are shown ± standard deviation. **(C)** Western blot analysis of *Yersinia* strains grown in low calcium LB at 26°C or 37°C for 3 hours. Equal amounts of cell lysates were probed for RpoA, IscR, H-NS, and YmoA as indicated. One representative experiment out of three biological replicates is shown. Statistical analysis was performed using an unpaired Student's t-test (n.s. non-significant).

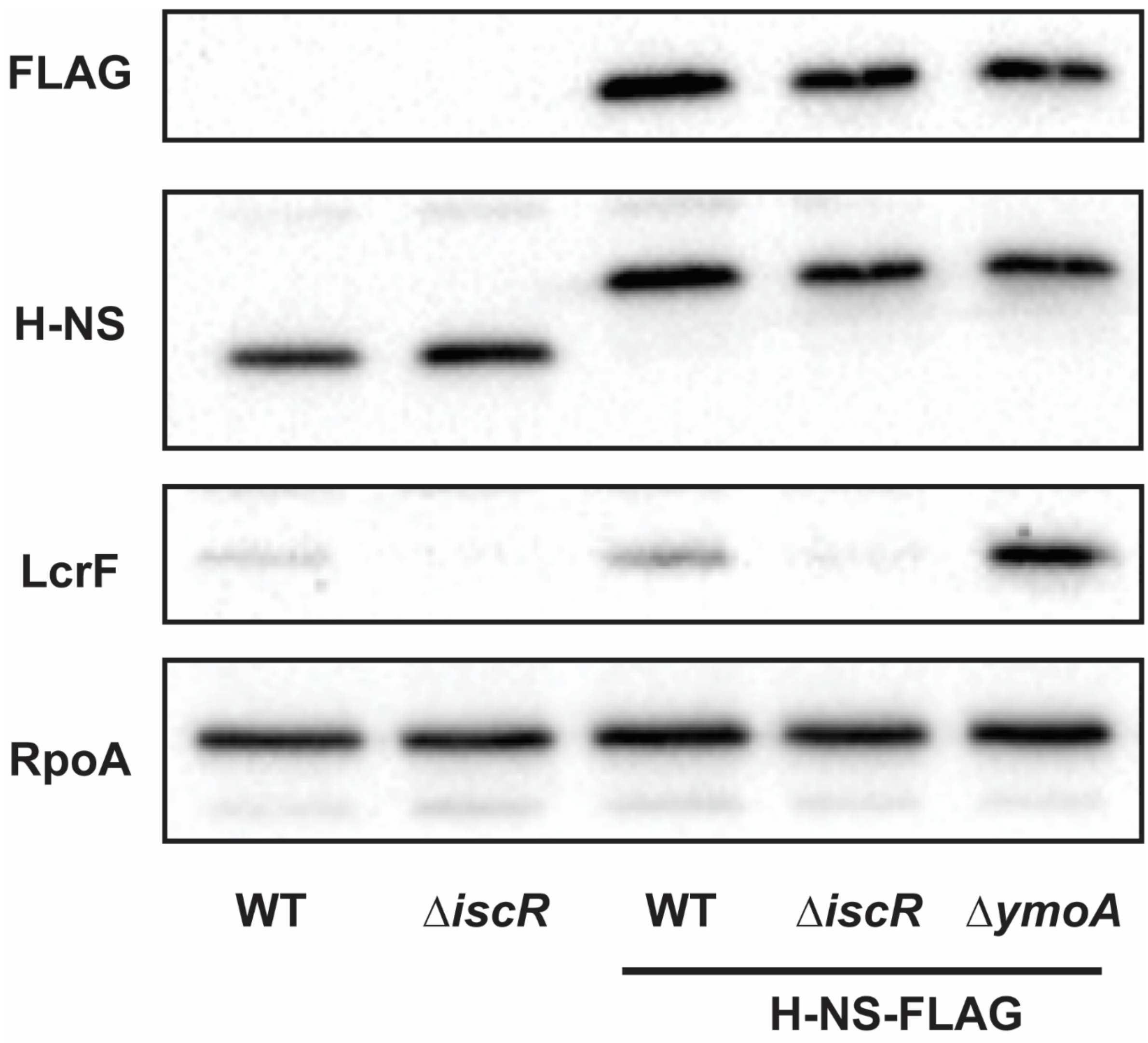


**Figure S4. 3xFLAG tag allows for detection of H-NS using FLAG antibody and does not affect H-NS ability to repress LcrF.** Western blot analysis of whole cell extracts from WT *Y. pseudotuberculosis* or a strain harboring a chromosomally-encoded 3xFLAG tagged H-NS visualized using anti-FLAG, anti-HNS, anti-LcrF, or anti-RpoA antibodies. One representative experiment out of three biological replicates is shown.

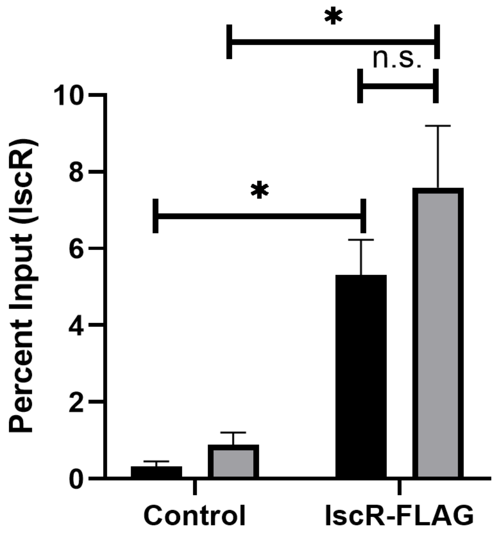


**Figure S5. IscR enrichment at the *suf* promoter is not influenced by temperature.** The relative enrichment (percent input) of *suf* promoter DNA analyzed by ChIP-qPCR with the IscR-FLAG strain or the control strain (WT; non-FLAG tagged IscR). ChIP-qPCR was performed with bacteria grown at 26°C (black bars) or 37°C (grey bars). The average of three independent replicates ± standard deviation is shown and statistical analysis was performed using an unpaired Student's t-test. (*p<.05, and n.s. non-significant).

**Table S1. Strains used in this study.**

| Strain | Relevant Genotype | Source or References |
|---|---|---|
| IP2666/(WT) | Naturally lacks full-length YopT | (1) |
| IP2666/(Δ*iscR*) | *iscR* in frame deletion of codons 2 to 156 | (2) |
| IP2666/(IscR 3xFLAG) | In frame C-terminus 3xFLAG tag of chromosomal IscR | (3) |
| IP2666/(H-NS 3xFLAG) | In frame C-terminus 3xFLAG tag of chromosomal H-NS | This work |
| IP2666/(Δ*ymoA*) | *ymoA* in frame full deletion | (4) |
| IP2666/(Δ*iscR*Δ*ymoA*) | Double deletion mutant of *iscR* and *ymoA* | This work |
| IP2666/( *ymoA*$^{D43N}$) | Single residue mutation of D43N YmoA | This work |
| IP2666/(Δ*iscR* *ymoA*$^{D43N}$) | *iscR* in frame deletion in YmoA D43N mutant | This work |
| IP2666/(lcrF$^{pNull}$) | Point mutations in IscR binding site upstream *yscW-lcrF* | (5) |
| IP2666/(Δ*ymoA* lcrF$^{pNull}$) | *ymoA* in frame deletion in lcrF$^{pNull}$ mutant | This work |
| IP2666/(Δ*lcrF*) | *lcrF* in frame full deletion | (6) |
| IP2666/(Δ*lcrF*Δ*ymoA*) | *ymoA* in frame deletion in *lcrF* mutant | This work |

**Table S2. *Y. pseudotuberculosis* primers used in this study.**

| Name | Primer Sequence[a] | References |
|---|---|---|
| qPCR_*16s*_F | AGCCAGCGGACCACATAAAG | (7) |
| qPCR_*16s*_R | AGTTGCAGACTCCAATCCGG | (7) |
| qPCR_*lcrF*_F | GGAGTGATTTTCCGTCAGTA | (2) |
| qPCR_*lcrF*_R | CTCCATAAATTTTTGCAACC | (2) |
| qPCR_*iscR*_F | CAGGGCGGAAATCGCTGCCT | (5) |
| qPCR_*iscR*_R | ATTAGCCGTTGCGGCGCCTAT | (5) |
| qPCR_*hns*_F | TGCAACAATACCGTGAAATG | This work |
| qPCR_*hns*_R | AGCACGTTTTGCTTTACCAG | This work |
| qPCR_*gyrA*_F | GGGGAAGTGGTGCTGAATAA | (8) |
| qPCR_*gyrA*_R | AAAATGGTACGGCGAGTCAC | (8) |
| qPCR_*cpxR*_F | TTGATGATGACCGTGAACTG | This work |
| qPCR_*cpxR*_R | ATCATAGGCGACCACAACAT | This work |
| qPCR_*rcsB*_F | GCAAATTGAATGGGTAAACG | This work |
| qPCR_*rcsB*_R | TAATTAGCACGTTGGCATCA | This work |
| qPCR_*ymoA*_F | CCTGATGCGTTTAAGAAAATG | This work |
| qPCR_*ymoA*_R | GATGGTCTGCAGCTGAGTAAA | This work |
| F*hns*_cds | cgaattcctgcagcccggggAGATGAGCACCATAAATG | This work |
| R*hns*_cds | aaccccccatCAACAGGAAGTCATCCAG | This work |
| F3xFLAG | cttcctgttgATGGGGGGTTCTGACTAC | This work |
| R3xFLAG | aataaaactaTCAACCTTTATCGTCGTCATC | This work |
| F3'*hns* | taaaggttgaTAGTTTTATTTCTTTAGCTATTACTATCG | This work |
| R3'*hns* | agggaacaaaagctggagctCGCCTAAATAGTCGTGGG | This work |
| F5'Δ*ymoA* | cgaattcctgcagcccggggGATAGACAGCTGTATTTATATGAC | This work |
| R5'Δ*ymoA* | gcgctaagcaGGTTTTTCTTCTCGATATACAAATTAATATTG | This work |
| F3'Δ*ymoA* | aagaaaaaccTGCTTAGCGCTGGTTAAG | This work |
| R3'Δ*ymoA* | agggaacaaaagctggagctCCTGTATTATCACTTTCCTGC | This work |
| pUC19-YmoA_F | acggccagtgaattcgagct*c*TTCATTTGTGATGAGTTTTAAAATAAAA TAC | This work |
| pUC19-YmoA-R | cctgcaggtcgactctagag*gatcc*AGAGGGCTGAATTTGAATG | This work |
| *ymoA*$^{D43N}$ _F | CTCAGCTGCAaACCATCGCCT | This work |
| *ymoA*$^{D43N}$ _R | TAAAACAATTCCAGTTCATCATCAGAAAGTTCG | This work |
| pFU99a_*ymoA*_F | cctttcgtcttcacctcgagTAATTGGTATATTTTCAATGCTTGTTTGGA TATCAATAC | This work |
| pFU99a_*ymoA*_R | ttcatttttaattcctcctgGTCATGCCGCTTAGGCGAG | This work |
| pFU99a_*hns*_F | cctttcgtcttcacctcgagATTGTACATAACGATACAGAAAC | This work |
| pFU99a_*hns*_R | ttcatttttaattcctcctgGTTGTTAAGAATTTTTAACGCTTC | This work |
| *hns*_gRNA_F | GCACTCCTAGTCTCAAATTATAAT | This work |
| *hns*_gRNA_R | AAACATTATAATTTGAGACTAGGA | This work |
| *hns*ChIP_site1_F | GCCCGTGCTCTTTATTGGG | This work |
| *hns*ChIP_site1_R | CACTTCAGCTGTGGCCTCTA | This work |
| *hns*ChIP_site2_F | TGGGGTGATTAACACCGG | This work |
| *hns*ChIP_site2_R | ATATAAGTGAACCTCTTGTTGGTTAAC | This work |
| *hns*ChIP_site3_F | TTATATGCGCAAGGTGTGATATTG | This work |
| *hns*ChIP_site3_R | TTCCCAATTATCTCAACGGGT | This work |
| *hns*ChIP_control_F | TGACGTCGGCAGTC | This work |
| *hns*ChIP_control_R | TCACCCCTTCGCAATAC | This work |
| *iscR*ChIP_*lcrF*_F | CGATATGGTTAACCAACAAGAGGTTC | This work |
| *iscR*ChIP_*lcrF*_R | GCACAGGAGAAATACAATTACCATAC | This work |
| *iscR*ChIP_*suf*_F | CTTTTAGACCTCCTTGGGTATCGC | This work |
| *iscR*ChIP_*suf*_R | CCGTTTGTTTTGCAGGGATATTAGG | This work |
| *iscR*ChIP_*hpt*_F | GCATGATGCTGGGCTTTAC | This work |
| *iscR*ChIP_*hpt*_R | ATAACAAAAATGCGCAGTGG | This work |
| pFU99a_*yscW*/*lcrF*_R | ttcatttttaattcctcctgAGAAATGATGAGTGCTATAATACG | This work |

| | | |
|---|---|---|
| pFU99a_*yscWlcrF*_p1 | cctttcgtcttcacctcgagCAAGTTCAGACTGTGCGC | This work |
| pFU99a_*yscWlcrF*_p2 | cctttcgtcttcacctcgagAGGCTGCAATGTAACTAG | This work |
| pFU99a_*yscWlcrF*_p3 | cctttcgtcttcacctcgagATGGTTAACCAACAAGAGG | This work |
| pFU99a_*yscWlcrF*_p4 | cctttcgtcttcacctcgagAATTAGGATTAATCTCTTGACTTTTTTTTG | This work |
| pFU99a_*yscWlcrF*_p5 | cctttcgtcttcacctcgagGGCTTTATATGCGCAAGG | This work |

[a] Uppercase specifies primer that anneals to target for molecular cloning, lowercase is complementary sequence for NEB Gibson Assembly or extra nucleotides to facilitate efficient restriction digest

**Table S3. Plasmids used in this study.**

| Name | Description | References |
|---|---|---|
| pPK7179-*yscW-lcrF* | Promoter template for *in vitro* transcription, $Amp^R$ | (3) |
| pPK7179-*sufA*-YP | Promoter template for *in vitro* transcription, $Amp^R$ | (3) |
| pPK7179-*sufA-EC* | Promoter template for *in vitro* transcription, $Amp^R$ | (9) |
| pSR47S Δ*ymoA* | Suicide vector for *ymoA* deletion, $Kan^R$ | This work |
| pUC19 YmoA | Vector with CDS of YmoA, $Amp^R$ | This work |
| pUC19 $ymoA^{D43N}$ | Vector with CDS of YmoA with $ymoA^{D43N}$ mutation, $Amp^R$ | This work |
| pSR47S $ymoA^{D43N}$ | Suicide vector for $ymoA^{D43N}$ mutation, $Kan^R$ | This work |
| pdCas9-bacteria | Expressing dCas9 protein under the control of an ATc-inducible promoter with a TetR casette, $Cm^R$ | (10) |
| pgRNA-tetO-JTetR | Expressing TetR driven by promoter J23119 and sgRNA driven by PL2tetO. $Amp^R$ | (11) |
| pgRNA-tetO-JTetR-H-NS | 20-bp targeting sequence for *hns* gene was inserted into pgRNA-tetO-JTetR, $Amp^R$ | This work |
| pFU99a | Empty vector carrying promoter-less *lacZ* fusion, $Cm^R$ | This work |
| pFU99a p*ymoBA*::*lacZ* | The promoter of *ymoBA* fused to *lacZ*, $Cm^R$ | This work |
| pFU99a p*hns*::*lacZ* | The promoter of *hns* fused to *lacZ*, $Cm^R$ | This work |
| pFU99a p*yscW-lcrF*::*lacZ* p1 | The promoter of *yscW-lcrF* (-505 - +294) fused to *lacZ*, $Cm^R$ | This work |
| pFU99a p*yscW-lcrF*::*lacZ* p2 | The promoter of *yscW-lcrF* (-309 - +294) fused to *lacZ*, $Cm^R$ | This work |
| pFU99a p*yscW-lcrF*::*lacZ* p3 | The promoter of *yscW-lcrF* (-166 - +294) fused to *lacZ*, $Cm^R$ | This work |
| pFU99a p*yscW-lcrF*::*lacZ* p4 | The promoter of *yscW-lcrF* (-47 - +294) fused to *lacZ*, $Cm^R$ | This work |
| pFU99a p*yscW-lcrF*::*lacZ* p5 | The promoter of *yscW-lcrF* (+101 - +294) fused to *lacZ*, $Cm^R$ | This work |

Yersinia pestis by using an optimized CRISPR interference system. Appl Environ Microbiol. 2019;
